# ER reorganization and intracellular retention of CD58 are functionally independent properties of the human cytomegalovirus ER resident glycoprotein UL148

**DOI:** 10.1101/746438

**Authors:** Christopher C. Nguyen, Anthony J. Domma, Hongbo Zhang, Jeremy P. Kamil

## Abstract

The human cytomegalovirus (HCMV) endoplasmic reticulum (ER)-resident glycoprotein UL148 is posited to play roles in immune evasion and regulation of viral cell tropism. UL148 prevents cell surface presentation of the immune cell costimulatory ligand CD58 while promoting maturation and virion incorporation of glycoprotein O, a receptor binding subunit for an envelope glycoprotein complex involved in entry. Meanwhile, UL148 activates the unfolded protein response (UPR) and causes large-scale reorganization of the ER. In an effort to determine whether the seemingly disparate effects of UL148 are related or discrete, we generated charged-cluster-to-alanine (CCTA) mutants of six charged clusters within the UL148 ectodomain, and compared them against wildtype UL148, in the context of recombinant viruses and in ectopic expression, assaying for effects on ER remodeling and CD58 surface presentation. Two mutants, targeting charged clusters spanning residues 79-83 (CC3) and 133-136 (CC4), respectively, retained the potential to impede CD58 presentation, and did so to an extent comparable to wildtype. Of the six mutants, only CC3 retained the capacity to reorganize the ER, showing a partial phenotype. Wildtype UL148 accumulates in a detergent-insoluble form during infection. However, all six CCTA mutants were fully soluble, which may imply a relationship between insolubility and organelle remodeling. Additionally, we found that the chimpanzee cytomegalovirus UL148 homolog suppresses CD58 presentation but fails to reorganize the ER, while the homolog from rhesus cytomegalovirus shows neither activity. Collectively, our findings illustrate varying degrees of functional divergence between homologous primate cytomegalovirus immunevasins and suggest that ER reorganization is unique to HCMV UL148.

**IMPORTANCE:** In myriad examples, viral gene products cause striking effects on cells, such as activation of stress responses. It can be challenging to decipher how such effects contribute to the biological roles of the proteins. The HCMV glycoprotein UL148 retains CD58 within the ER, thereby preventing it from reaching the cell surface where it functions to stimulate cell-mediated antiviral responses. Intriguingly, UL148 also triggers the formation of large, ER-derived membranous structures, and activates the UPR, a set of signaling pathways involved in adaptation to ER stress. We demonstrate that the potential of UL148 to reorganize the ER and to retain CD58 are separable by mutagenesis and possibly, by evolution, since chimpanzee cytomegalovirus UL148 retains CD58 but does not remodel the ER. Our findings imply that ER reorganization contributes to other roles of UL148, such as modulation of alternative viral glycoprotein complexes that govern the virus’ ability to infect different cell types.

## INTRODUCTION

Cytomegaloviruses (CMVs), like other herpesviruses, encode a large and multifarious array of gene products and regulatory RNAs that alter host cell functions to support viral replication, spread, and lifelong persistence [reviewed in (1, 2)]. CMVs have evolved a number of functions that dampen antiviral responses that otherwise produce sterilizing immunity. Rodent and primate cytomegalovirus (CMV) species, such as mouse cytomegalovirus (MCMV) and human cytomegalovirus (HCMV), express a large number of proteins and microRNAs that interfere with antigen presentation and prevent activation of cytotoxic T cells and NK cells [reviewed in (3–6)], (7). In HCMV, several viral gene products involved in immune evasion, such as UL141 (8, 9), UL142 (10–12), and UL148 (13), are encoded within a region of the genome termed the ‘UL*b’*,’ which undergoes extensive rearrangements and deletions during extensive serial passage in cultured cells (14–16).

The HCMV glycoprotein UL148 impedes the activation of cytotoxic T-cell and NK cell mediated responses by binding the co-stimulatory ligand CD58 and retaining it within the endoplasmic reticulum (ER) (13). Intriguingly, UL148 also shows a strong capacity to influence HCMV cell tropism, with *UL148*-null mutant viruses showing ∼100-fold improved replication in epithelial cells (17, 18). The evidence suggests that UL148 stabilizes immature forms of glycoprotein O (gO) within the ER, thereby promoting the maturation and incorporation into virions of the heterotrimeric glycoprotein H (gH)/ glycoprotein L (gL)/ gO complex. gH/gL/gO (Trimer) and gH/gL/UL128-131 (Pentamer) are the two alternative HCMV gH/gL complexes that endow the virus with the ability to utilize different proteinaceous host cell surface factors as entry receptors [reviewed in (19)]. In particular, gO, in the context of Trimer, is required for viral entry via the platelet-derived growth factor receptor alpha (PDGFR*α*), which is well expressed in fibroblasts but poorly if at all in epithelial cells (20–23). Infection of epithelial and endothelial cells, on the other hand, appears to require direct interactions between the Pentamer and neuropilin-2 (Nrp2) (24), together with roles for OR14I1 (25) and CD147 (26), addition to roles for the Trimer (27, 28). The striking replication advantage of *UL148*-null viruses in epithelial cells is presumably explained by decreased expression of Trimer, which in turn is thought to favor Pentamer-dependent interactions that are important for infection of epithelial cells.

Furthermore, UL148 is necessary and sufficient to activate the unfolded protein response (UPR) (29), a canonical ER-stress adaptation pathway conserved among eukaryotes. Although the mechanism for UPR activation remains unknown, UL148 appears to interact with the core ER-associated degradation (ERAD) factor SEL1L (18). Hence, UL148 might be hypothesized to cause ER stress by impairing the ability of the SEL1L/Hrd1 complex to dispose of misfolded proteins. Furthermore, UL148 drastically reorganizes the ER, causing the accumulation of unusual electron-dense membranous structures that appear to derive from distended rough-ER cisternae (30). These ER structures are replete with factors involved in glycoprotein quality control such as SEL1L, Hrd1, and EDEM1. We wondered whether the dramatic effects of UL148 on the ER might be related to its potential to retain CD58. We therefore sought to develop experimental approaches to interrogate whether these two UL148-dependent activities were separable or interdependent.

In the absence of a three-dimensional structure, charged-cluster-to-alanine (CCTA) mutagenesis provides a useful approach to identify functionally important residues within a protein sequence. The underlying rationale is based on the assumption that charged clusters of amino acids are more likely to be found on solvent exposed surfaces in the context of the tertiary structure of a given polypeptide, and therefore mutations affecting such residues are less likely to strongly influence overall protein folding or stability (31–34). Additionally, charged clusters are often involved in biologically relevant protein-protein interactions, as evidenced by the historical utility of CCTA mutagenesis for identifying residues involved in ligand-receptor interactions (31, 35–38). To define which UL148-associated activities may arise from shared versus distinct molecular mechanisms, we combined CCTA mutagenesis with marker-less bacterial artificial chromosome (BAC) recombination (“recombineering”) methods (39, 40) to generate HCMV genomes mutated for individual charged clusters within *UL148*. We also constructed vectors for transient ectopic expression of UL148 CCTA mutants in non-infected cells. Finally, we cloned UL148 homologs from other primate cytomegaloviruses, including chimpanzee CMV (53% amino acid identity with UL148) and rhesus macaque CMV (25% amino acid identity), so as to gain insight into potential evolutionary relationships between the various activities of HCMV UL148.

## RESULTS

### UL148 charged-cluster-to-alanine mutagenesis

In order to identify charged residues within UL148 in an unbiased manner, we utilized a simple scanning algorithm (window = 6 residues) to analyze the predicted ectodomain of UL148. We designated any window containing at least 40% charged amino acids (K, R, D, E) to be a charged cluster (**FIG 1A**). For practical purposes, we opted to treat as a single cluster three clusters that were found in very close proximity to each other (residues 220-231). In this manner, we were able to focus our mutagenesis strategy on six charged clusters within the UL148 ectodomain (**FIG 1B**). We constructed a series of expression plasmids encoding HA-tagged UL148 harboring CCTA mutations at each of the clusters. For comparative purposes, we also cloned UL148 homologs from two other primate CMVs—notably UL148 homologs are found only in primate CMVs. In particular, we constructed plasmids encoding chimpanzee CMV UL148 (Ch148; 53% identity) and rhesus CMV Rh159 (25% identity), each appended at its C-terminus to an influenza virus hemagglutinin (HA) epitope tag.

**FIGURE 1.**
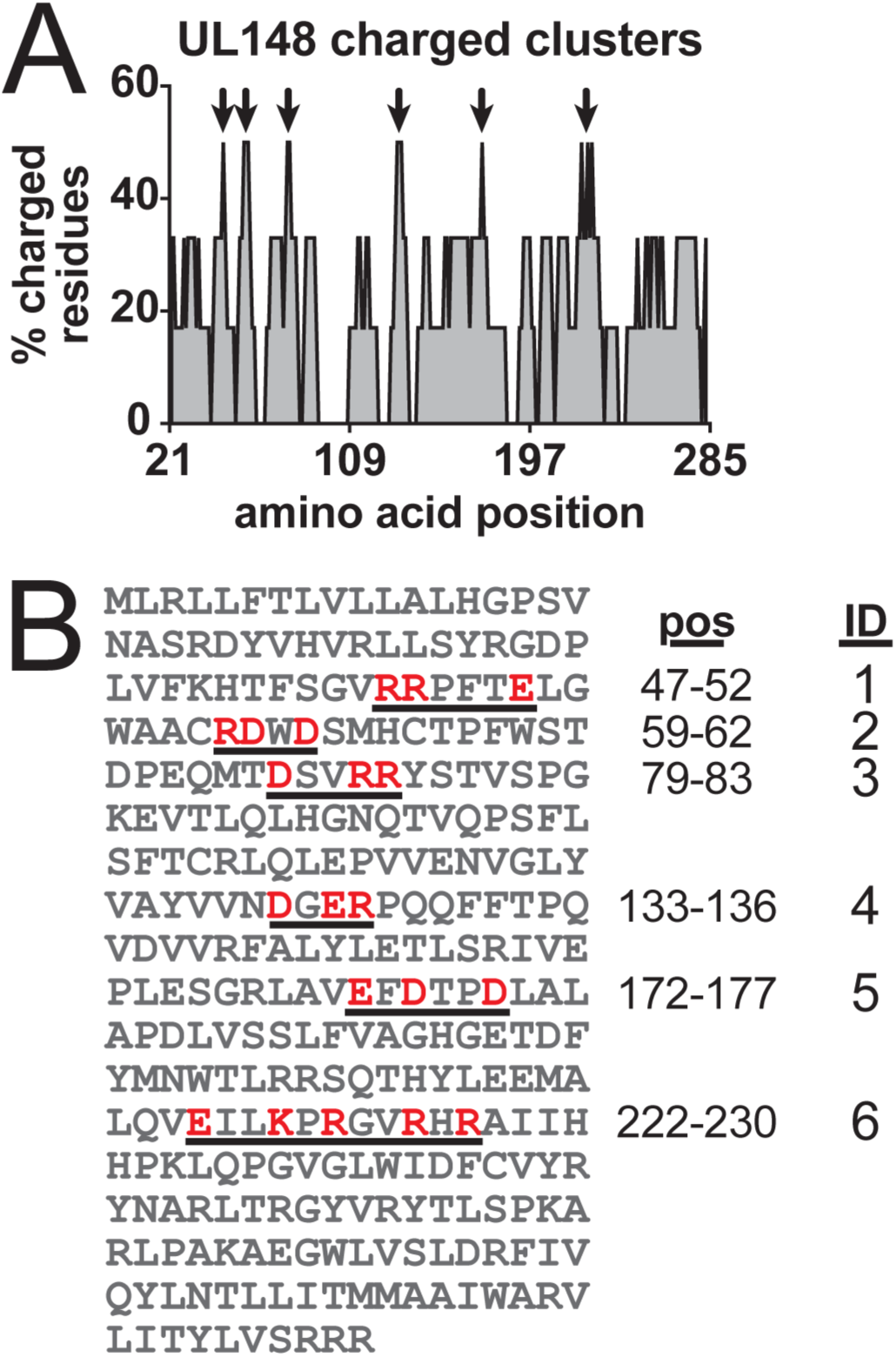
Charged amino acid clusters within the predicted UL148 ectodomain. (A) Percentage of charged residues (K, R, D, E) within the predicted UL148 ectodomain identified using an iterative scanning window six amino acid residues in size. Black arrows designate charged clusters (>40% charged residues). The last three local maxima in close proximity were treated as a single cluster. (B) UL148 protein sequence. Amino acid positions (pos) of the identified charged clusters are underlined. Residues in red were mutated to alanine in the six constructs indicated above using sequential numbering (ID), starting from the amino-terminus.

### Expression of UL148 CCTA mutants and UL148 homologs

As a first step to confirm the expression of the UL148 CCTA mutants, and the primate CMV homologs of UL148, we transfected HEK-293T (293T) cells with the different expression plasmids. Robust levels of protein expression were observed for both Rh159 and Ch148 (**FIG 2**). HCMV UL148 expressed to visibly lower levels than Rh159 or Ch148 during transient overexpression in transfected cells. We interpret the somewhat lower peak expression level of HCMV UL148 as likely being due to its activation of the unfolded protein response (29). Nonetheless, each of the CCTA mutants expressed at levels comparable to that of wildtype UL148, except perhaps CCTA mutant #6, which affected the largest cluster (residues 220-231). Overall, each of the CCTA constructs was readily detected following transient transfection of 293T cells (**FIG 2**).

**FIGURE 2.**
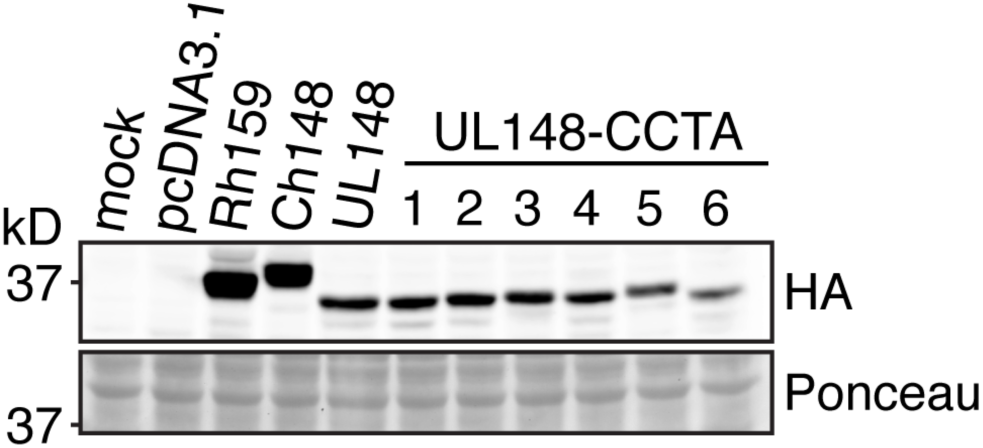
Expression of HA-tagged UL148 CCTA mutants and UL148 homologs. Human embryonic kidney 293T cells were transfected with expression plasmids encoding the indicated HA-tagged proteins. At 48 h post-transfection, cell lysates were analyzed by Western blot.

### Disruption of four out of six charged clusters abolishes UL148-dependent suppression of cell surface CD58

Wang *et al*. (13) reported that HCMV UL148 is necessary and sufficient to suppress surface presentation of CD58, a ligand for CD2 that co-stimulates T and NK cell activation, such as in response to HCMV infection. Since CCTA mutagenesis has been extensively used to define “interface” residues involved in protein-protein interactions, we asked whether UL148 mutants ablated for specific charged clusters would show differences in their ability to suppress CD58 surface presentation. Turning to a transient expression system to ask this question, we electroporated UL148-CCTA constructs into A549 lung adenocarcinoma cells, which basally express surface CD58 (see below). In order to limit our analyses to cells that took up plasmid, we co-transfected a GFP expression vector along with each UL148 construct. Gating for GFP positive cells, we observed a ∼40% decrease in CD58 median fluorescence intensity (MFI) in cells transfected with wildtype UL148 (**FIG 3A**). A comparable decrease in surface CD58 was detected in cells transfected with Ch148 expression vector but not with the vector expressing the rhesus CMV UL148 homolog, Rh159. Importantly, four of our six CCTA mutants failed to downregulate surface CD58. The remaining two CCTA constructs, however, mutated at charged clusters #3 or #4, (**FIG 1B**), suppressed CD58 surface presentation comparably to wildtype UL148.

**FIGURE 3.**
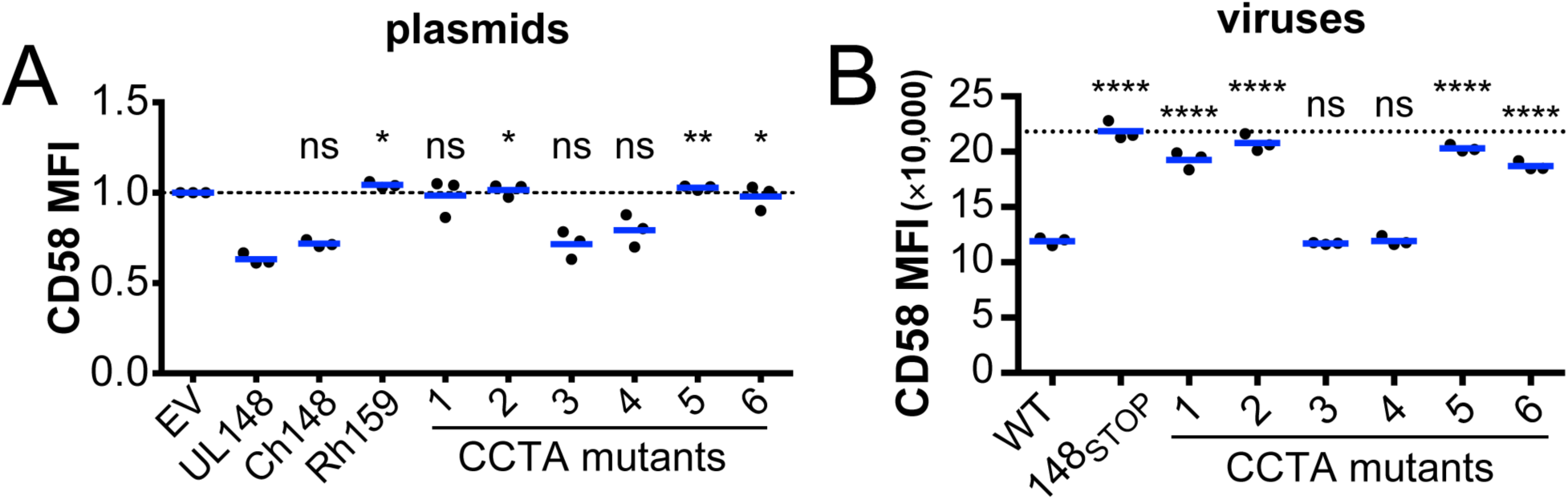
Analysis of cell surface CD58 levels during expression of UL148 CCTA mutants and homologs from other primate CMVs. (A) A549 cells were transfected with plasmids expressing the indicated protein or empty vector (EV). At 72 h post-transfection, cell surface CD58 levels were analyzed by flow cytometry. Median fluorescence intensity (MFI) values were normalized to the EV control. (B) Fibroblasts (HFFT) were infected at an MOI of 2 TCID_50_ per cell with HCMV strain TB40/E derivatives wildtype for *UL148* (WT), carrying the indicated UL148 CCTA mutations, or *UL148*-null (148_STOP_). At 72 h post-infection, cell surface CD58 was quantified by flow cytometry. (A-B) Data points indicate biological replicates from three independent experiments. P-values comparing the mean of each condition to wildtype UL148 (UL148, panel A) or wildtype virus (WT, panel B) were obtained by one-way analysis of variance (ANOVA) with Tukey’s correction for multiple comparisons: ns, not significant; *, p < 0.05; **, p < 0.01; ****, p < 0.0001.

We next sought to analyze the importance of the charged clusters in the context of infection. We employed *en passant* mutagenesis (39, 40) to construct HCMV recombinants harboring these CCTA mutations. Upon infection of fibroblasts with the CCTA mutant viruses, we obtained results that largely recapitulate our findings from electroporation of A549 cells with plasmid expression vectors (**FIG 3B**). The four different charged clusters identified to be required for CD58 suppression in transient transfection were likewise required to suppress CD58 surface presentation during HCMV infection. Moreover, the two CCTA mutants competent for CD58 suppression in transfected A549 cells, CC3 and CC4, likewise appeared to suppress CD58 and did so indistinguishably from parental wildtype virus. These results suggest that the two charged clusters at 79-83 (CC3) and 133-136 (CC4) are dispensable for intracellular retention of CD58. Additionally, our results indicate that the other four charged clusters are either directly required to prevent surface presentation of CD58, or otherwise necessary for UL148 to properly fold or function, since these mutant viruses phenocopy *UL148*-null virus (148_STOP_) in failing to downregulate CD58 surface levels.

### The propensity of UL148 to accumulate in a detergent-insoluble form is dispensable for suppression of CD58 surface presentation

We recently observed that UL148 accumulates in a detergent-insoluble form during infection (30). We therefore wished to examine whether any of the CCTA mutations affected the solubility of UL148 during infection, since this could provide a clue as to whether the UL148 insolubility is required for retention of CD58. We carried out these experiments using fibroblasts infected with our series of recombinant HCMVs expressing the six different CCTA mutants, so as to minimize the potential for artifacts concerning solubility that might confound results obtained from transient overexpression of plasmids. Each of the CCTA mutants failed to accumulate in the detergent-insoluble fraction, while wildtype UL148, as expected appeared in both the soluble and insoluble fractions (**FIG 4**). Nonetheless, cells infected with CCTA mutants affecting the clusters #3 (residues 79-83) and #4 (residues 133-136) showed defects in CD58 maturation matching those seen with wildtype virus (**FIG 4**), which suggests that the potential for UL148 to accumulate in a detergent-insoluble form is separable from its capacity to prevent CD58 maturation.

**FIGURE 4.**
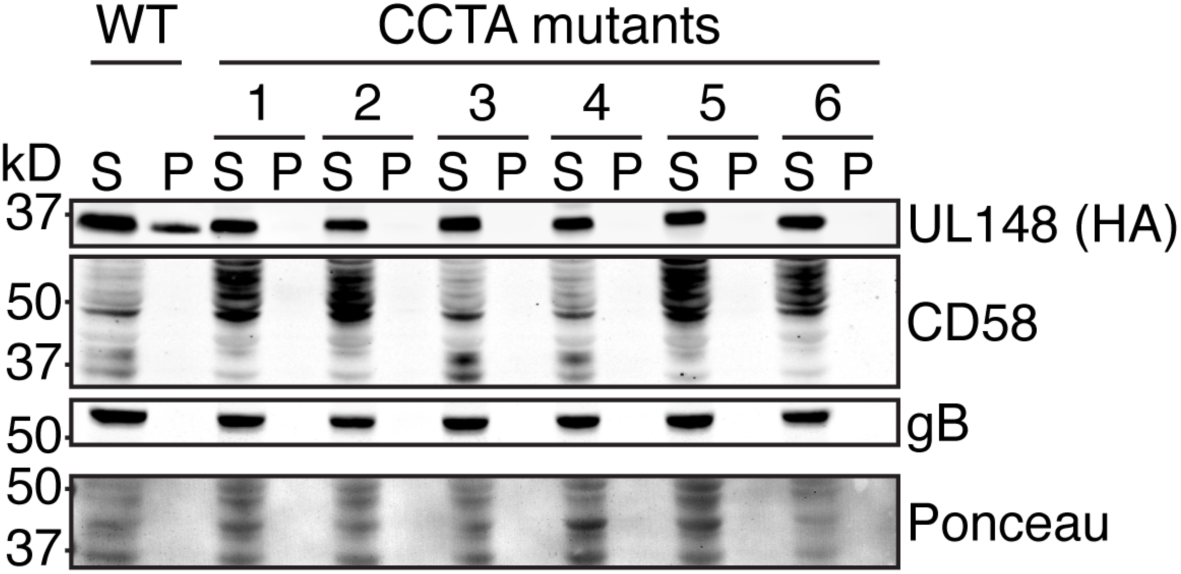
The propensity of UL148 to accumulate in a detergent-insoluble form is not required to block CD58 maturation. Fibroblasts (HFFT) were infected at an MOI of 1 TCID_50_ per cell with wildtype HCMV (WT) or the indicated *UL148* mutant (CCTA mutants 1-6). At 72 h post-infection, cell lysates were separated by centrifugation into supernatant (S) containing soluble proteins and pellet (P) containing insoluble proteins. Supernatant and pellet fractions were analyzed by Western blot for UL148, CD58, and gB levels.

### Five out of six charged clusters are necessary for UL148-mediated ER remodeling

UL148 activates the unfolded protein response (UPR), both during infection and ectopic expression (29), and is also necessary and sufficient to drastically remodel the ER (30). Hoping to identify charged clusters necessary for these striking effects, we again turned to our panel of CCTA mutants. Results from confocal immunofluorescence microscopy of infected cells stained for Hrd1 and UL148 (HA tag) at 72 h post-infection (hpi) indicated that five of the six CCTA mutants matched the phenotype seen for *UL148*-null virus (148_STOP_) in failing to cause ER remodeling, as indicated by Hrd1 staining (**FIG 5**). The remaining mutant (CC3) exhibited a partial phenotype wherein UL148 puncta formation could be detected in a minority of cells. Interestingly, the CC4 mutant completely lost the ability to remodel the ER but nonetheless blocks maturation and surface presentation of CD58 similarly if not equivalently to wildtype (**FIG 3-4**). Because both CC3 and CC4 mutants block CD58 surface presentation while exhibiting, respectively, either a significant decrease or a complete loss of ER remodeling, our results suggest that these two activities involve distinct molecular mechanisms.

**FIGURE 5.**
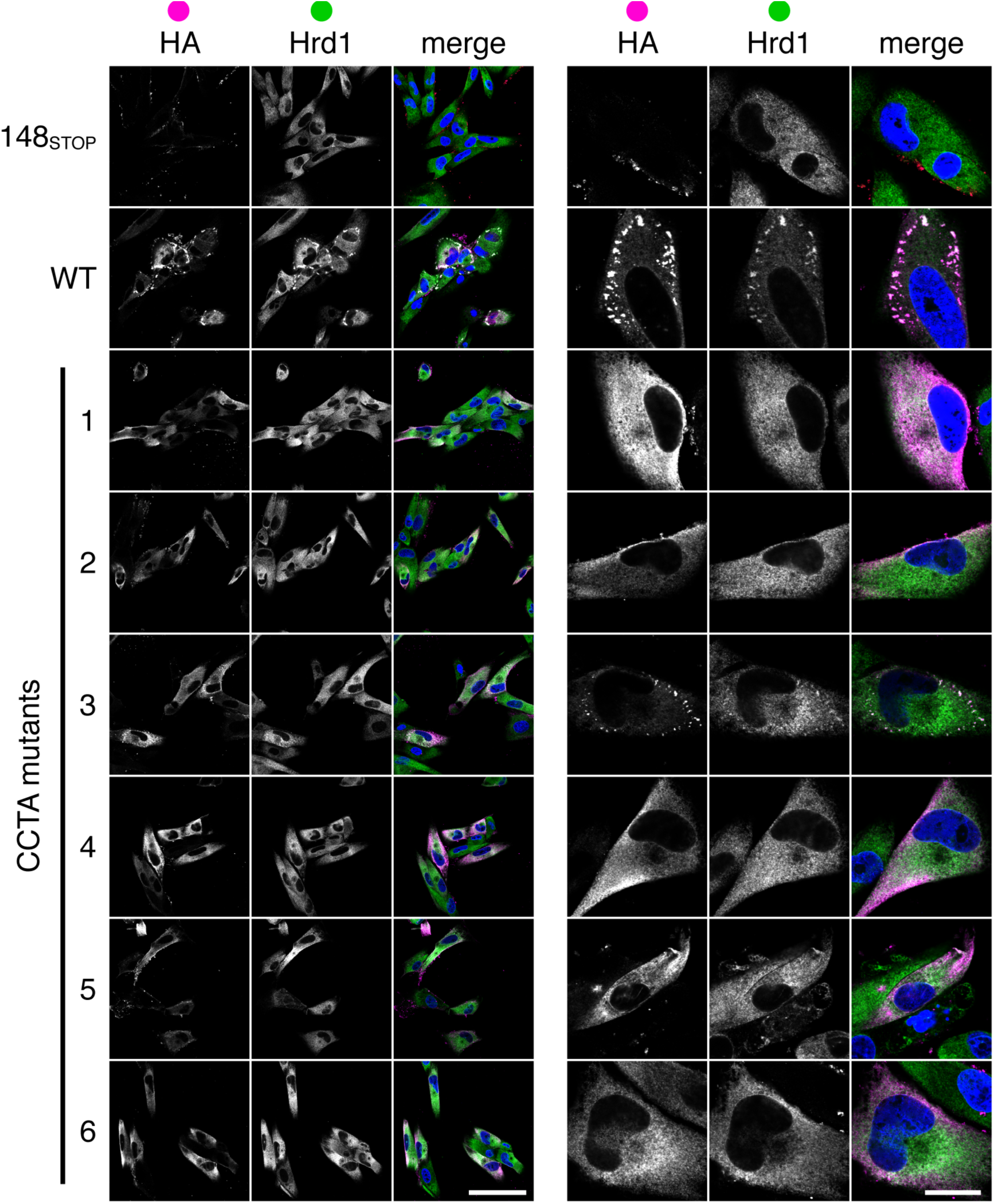
Mutation of five UL148 charged clusters abolishes punctate co-localization of UL148 and ER marker Hrd1. Fibroblasts (HFFT) were infected at an MOI of 1 TCID_50_ per cell with HCMV strain TB40/E wildtype for *UL148*, harboring the indicated *UL148* CCTA mutant, or entirely null for *UL148* (148_STOP_). At 72 h post-infection, cells were fixed and imaged by confocal microscopy after staining with antibodies specific for HA (UL148^HA^, magenta) or ER marker Hrd1 (green). Nuclei were counterstained using Hoechst dye (blue).

### The chimpanzee CMV UL148 homolog does not remodel the ER

Lastly, we sought to determine whether the effects of UL148 on the ER and on CD58 surface expression might be conserved by assaying HCMV UL148 against its homologs from other primate CMVs. Rh159, the rhesus CMV homolog, shares 25% amino acid identity with HCMV UL148 and has been shown to retain NKG2D receptor activating ligands of the MIC- and ULBP families (7). Furthermore, Rh159 fails to activate the UPR and does not appear to remodel ER membranes (29, 30). However, the chimpanzee CMV homolog of UL148, Ch148, shares 53% amino acid identity with UL148, and to the best of our knowledge has not been examined for its effects on CD58 or on the ER. We cloned Ch148 fused at its predicted C-terminal tail to enhanced GFP (GFP) in the context of an inducible expression vector to facilitate observation of Ch148 localization patterns within the cell. We previously constructed similar vectors encoding Rh159 and UL148 fused to GFP (30). Upon induction of protein expression using 100 ng/mL doxycycline, we observed the formation of punctate structures for UL148-GFP but diffuse, reticulate signal for Rh159-GFP, as expected (30). Interestingly, Ch148-GFP exhibited a diffuse localization pattern more similar to that of Rh159 than UL148 (**FIG 6**). Taken together with the results from our series of CCTA mutants, this result suggests that ER remodeling and CD58 retention arise from distinct mechanisms.

**FIGURE 6.**
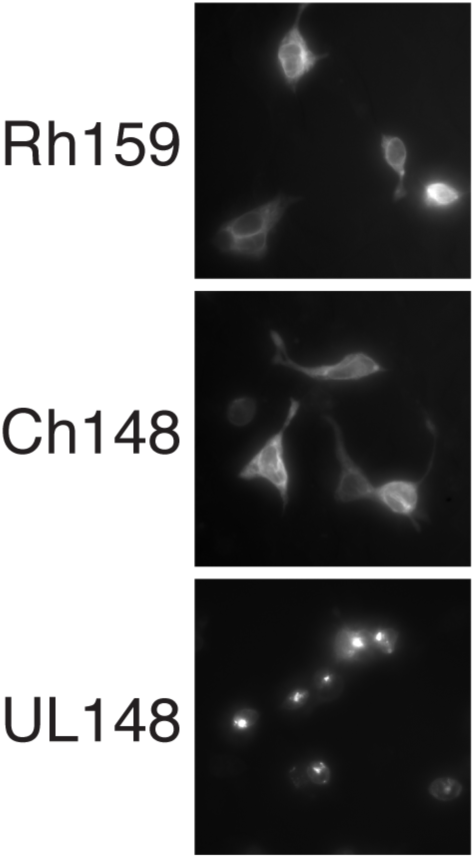
Localization of UL148, Rh159, and Ch148. Human embryonic kidney 293T cells were transfected with doxycycline-inducible vectors encoding the indicated protein fused to a GFP tag. 100 ng/mL doxycycline was added to cells 1 h post-transfection. At 48 h post-transfection, GFP signal was visualized by immunofluorescence microscopy.

## DISCUSSION

### Evidence of distinct mechanisms for CD58 suppression and ER remodeling

Upon discovering that UL148 reorganizes the ER (30), we wondered whether this phenotype might play a role in ER-retention of CD58. We considered the possibility that perhaps CD58 is so efficiently chaperoned to the cell surface that it somehow resists being sequestered via simple high affinity interactions with an ER-resident protein. So perhaps UL148 forms bizarre membranous structures to more potently obstruct its presentation at the cell surface? Two findings from this study argue against this possibility, and instead indicate that ER reorganization is dispensable for UL148-mediated interference with CD58 surface presentation. Firstly, the CCTA mutant affecting charged cluster #4 fails to remodel the ER but impedes CD58 surface presentation comparably, if not indistinguishably, from wildtype UL148 (**FIG 3-5**). Secondly, the UL148 homolog from chimpanzee CMV (Ch148) is sufficient to suppress surface CD58 presentation but fails to induce the formation of punctate structures that we consistently observe during expression of HCMV UL148, which indicate ER reorganization (30) (**FIG 3A, 6**). Although we cannot exclude the possibility that Ch148 suppresses CD58 surface presentation by a mechanism distinct from that of HCMV UL148, we consider this to be unlikely, considering the extensive sequence homology between Ch148 and HCMV UL148 (>50% amino acid identity). A summary view of the observed activities for Ch148 and UL148 constructs is shown in **FIG 7**. Collectively, the data suggest that the properties of UL148 that give rise to ER re-organization are distinct from those involved in intracellular retention of CD58.

**FIGURE 7.**
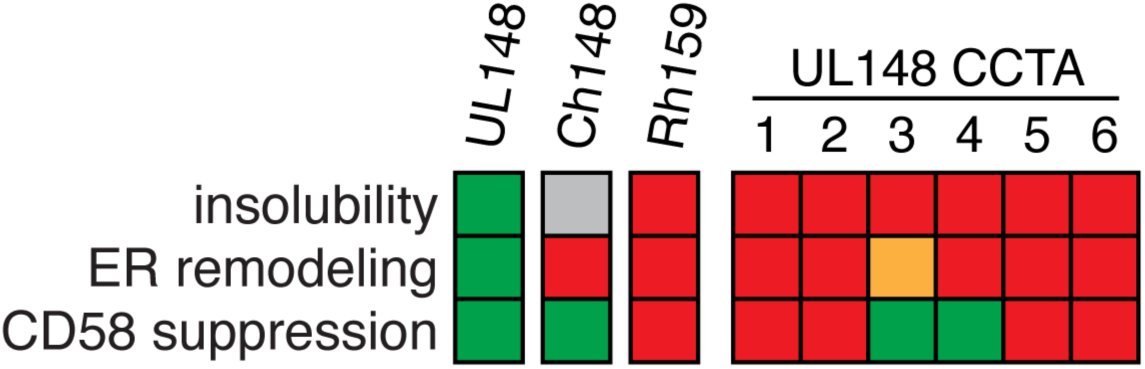
Summary of activities of CCTA mutants of UL148 and UL148 homologs. Green squares indicate activity indistinguishable from wild-type HCMV UL148. Red squares indicate total absence of the activity. Yellow squares indicate a partial phenotype. Gray square, not tested.

### Why does UL148 accumulate in a detergent-insoluble form, and what is the relationship to ER reorganization?

Fundamental to the biology of the ER is its system of mechanisms, such as N-glycosylation and ER-resident chaperones, that function to prevent proteins from aggregating or accumulating in an insoluble state (41, 42). Therefore, it seems unusual that the wildtype form of an ER resident glycoprotein like UL148 would accumulate in a detergent-insoluble form in the first place. Moreover, this feature of UL148 stands in stark contrast with every other glycoprotein we have analyzed in infected cells, including gB and CD58 (**FIG 4**), the RhCMV UL148 homolog Rh159 (30), as well as gH and gO (C.C.N. and J.P.K., unpublished results). Surprisingly, all six of the CCTA mutants examined in this study failed to accumulate in a detergent-insoluble form. Thus, a property readily disrupted by CCTA mutagenesis is required for the protein to accumulate in an insoluble form.

Notably, insoluble forms of many different proteins characteristically accumulate during treatments that diminish proteasomal or lysosomal protein degradation pathways (43–45). Conversely, protein aggregates are themselves thought to inhibit the ubiquitin proteasome system (46). Intriguingly, UL148, exhibits a marked tendency toward insolubility even in the absence of inhibitors affecting rates of protein degradation. We hypothesize that the native conformer adopted by wildtype UL148 may resist ER associated degradation (ERAD), and perhaps even generally inhibit ERAD. On the other hand, other examples of proteins that form ER structures, such as the Z-allele of alpha-1-antitrypsin (*SERPINA*) (47, 48), or the A103E mutant of the calcium channel subunit ORAI1 (*CRAMC1*), exhibit a propensity to aggregate or polymerize within the ER, which may make them resistant to ERAD. Given their improved solubility, each of the UL148 CCTA mutants, in contrast, might be predicted to show improved susceptibility to ERAD, and may also indicate that these mutants no longer aggregate or polymerize.

Strikingly, five of the six CCTA mutants fully lost the ability to reorganize the ER, while the one mutant that still showed ER organization exhibited a partial phenotype. It is notable that in all cases, the loss of ER reorganization correlated with markedly improved solubility. These observations collectively suggest that the propensity of UL148 to accumulate in an insoluble form may be mechanistically important for ER remodeling.

### Roles of ER remodeling cell tropism versus roles in immune evasion?

Collectively, our results suggest that ER remodeling is indicative of, if not directly involved, in a function unrelated to CD58 retention, since ER remodeling appears to be dispensable for this function (**FIG 3-6**). Accordingly, one might hypothesize ER reorganization to play roles in the cell tropism effects of UL148. The effects of UL148 on HCMV cell tropism correlate with its potential to promote the expression of gH/gL/gO, a heterotrimeric complex that endows virions with the ability to utilize PDGFR*α* as an entry receptor (20, 21, 49, 50). We previously found that gO behaves as a constitutive ERAD substrate and that UL148 stabilizes its expression (18). Given that the ER structures induced by UL148 contain huge amounts of glycoprotein quality control proteins that participate in ERAD (30), it seems plausible that UL148 may limit degradation of immature gO by sequestering the ERAD machinery. Despite that a role for ER remodeling in suppression of CD58 surface presentation seems unlikely, given our findings here, a role in regulation of glycoprotein complexes involved in cell tropism remains to be disproven.

### Proper folding of UL148 harboring CCTA mutations

In the absence of characterized monoclonal antibodies against UL148, we are limited in our ability to conclusively demonstrate that the UL148 species carrying mutated charged clusters fold properly in the ER. That said, our results offer some insight into the overall stability of the proteins expressed from our plasmid constructs and recombinant viruses. It is notable that all of six of the CCTA mutants we examined in this study show improved solubility relative to wildtype UL148. Insolubility is often associated with misfolding and aggregation. Therefore, the observation that wildtype UL148 is less soluble that its CCTA mutants may suggest that insolubility is important for one or more of its biological functions, but not for CD58 retention, as our results have identified two CCTA mutants that retain CD58 without accumulating in an insoluble form. Even so, caution compels us to avoid making strong conclusions from the other four CCTA mutants that have so far only recapitulated *UL148*-null phenotypes with regard to ER remodeling and CD58 retention. We are therefore focusing on mutants of charged clusters #3 (CC3) and #4 (CC4) in our ongoing studies concerning UL148-mediated ER remodeling, since CCTA mutants of these clusters can be presumed to fold properly, given that they remain able to suppress CD58 maturation.

### Inferences into the UL148-CD58 interaction face

Alanine scanning and CCTA mutagenesis approaches have proven useful to define the specific amino acid residues underlying biologically relevant protein-protein interactions. Despite that detailed empirical data on the tertiary structure of UL148 as-of-yet remain unavailable, the results from our studies of UL148 CCTA mutants support a number of inferences concerning the physical basis for its interaction with CD58. The two charged clusters that do not appear to be important for CD58 retention, #3 and #4, are found in a contiguous stretch of amino acids (residues 79-136) that lacks other charged clusters. Assuming our other four CCTA mutants do not grossly compromise UL148 folding, residues from the N-terminal half (residues 47-62) and C-terminal half (residues 172-230) of the UL148 ectodomain may participate in an interaction face with CD58. Likewise, our data suggest that the contiguous sequence containing clusters #3 and #4 represents a region within the 3D structure that does not participate in CD58 binding.

### Chimpanzee CMV provides a valuable model for comparative studies of HCMV gene products in cell-based assays

Taken together, our findings highlight the various functionally divergent roles among three different UL148 homologs in CMVs endemic to humans, rhesus macaques, and chimpanzees. While all three proteins act as ER-resident immunevasins, the rhesus CMV homolog lacks the potential to trigger the UPR (29, 30), and apparently functions to evade NK cells by preventing MIC- and ULBP family NKG2D ligands from reaching the cell surface (7). Chimpanzee and human CMV UL148 target CD58 instead of NKG2D ligands [(13), **FIG 3**)], but only HCMV UL148 is capable of remodeling the ER (**FIG 6**), a property that appears to depend on the capacity of UL148 to activate the integrated stress response downstream of UPR activation (30). Although chimpanzee CMV is more closely related to HCMV than rhesus CMV (51), the use of chimpanzees as an animal model for human diseases is prohibited, which likely has limited the interest of researchers in studying the virus. Our study, however, highlights the utility of studying chimpanzee CMV gene products, even in the context of cultured cells, to gain comparative insights into the functions of HCMV genes. Chimpanzee CMV homologs may be of particular importance for helping us to understand the functions of genes in regions of the virus such as the UL*b’*, which are not well conserved, if at all, in rodent CMVs. Moreover, at least in the example of UL148, the chimpanzee homolog appears to be more functionally aligned with HCMV UL148 than the rhesus CMV homolog. Because genes within the UL*b’* play crucial roles in latency, persistence, cell tropism, and immune evasion, elucidating their functions will be of critical importance. A clear picture of which gene specific functions are shared and divergent between the different primate CMVs may provide profound insights into the unique biology of HCMV.

## MATERIALS AND METHODS

### Cells

Human telomerase (hTERT)-immortalized human foreskin fibroblasts (HFFT) were described previously (18), and were cultured in Dulbecco’s modified Eagle’s medium (DMEM; Corning Cat #10-013-CV) supplemented with 25 µg/mL gentamicin sulfate (Invitrogen), 10 µg/mL ciprofloxacin HCl (Genhunter, Inc.) and 5% newborn calf serum (NCS) (Sigma #N4637). Human embryonic kidney 293T (HEK-293T) cells were obtained from Genhunter, Inc. (Nashville, TN), and cultured in DMEM supplemented as above except with 10% fetal bovine serum (FBS) (MilliporeSigma Cat #F2442) instead of NCS. A549 adenocarcinoma cells were obtained from ATCC (Cat # CCL-185) and grown in Ham’s F-12K medium (ThermoFisher, Cat # 21127022) supplemented with 10% FBS and antibiotics, as above.

### Virus

Virus was reconstituted by electroporation of purified bacterial artificial chromosome (BAC) DNA into HFFT and amplified until 100% cytopathic effect (CPE) was observed, as previously described (17, 18, 52). Cell-free virus in the medium was combined with cell-associated virus, which was released by Dounce-homogenization of the cells. Virus preparation was clarified of cellular debris by centrifugation (4000 × *g*, 10 min), followed by pelleting of virus by ultracentrifugation (85,000 × *g*, 1 h, 4°C) through a 20% sorbitol cushion. The supernatant was removed, and the viral pellet was resuspended in DMEM supplemented with 20% NCS and stored at -80°C.

### Virus titration

Infectivity of virus stocks was determined according to the tissue culture infectious dose 50% (TCID_50_) method. In brief, stocks were serially diluted, and diluted virus was used to infect eight wells per dilution in a 96-well plate seeded one day prior with 3 × 10^4^ HFFT per well. Infection was allowed to proceed for 9 days before IE1 staining as follows. Cells were washed (3 × 200 µL PBS per well) and fixed in ice-cold methanol (200 µL/well, 15 min at -20°C). Cells were washed (3 × 200 µL PBS per well) and stained with 1:200 mouse anti-IE1 primary antibody (supernatant from IE1 mAb 1B12 hybridoma cells, a generous gift of Thomas Shenk, Princeton University) for 1 h at 37°C. Cells were washed (3 × 200 µL PBS per well) and stained with 1:1000 goat anti-mouse Alexa Fluor 488 secondary antibody (ThermoFisher, Cat # A-11001) for 1 h at 37°C. The cells were washed (1 × 200 µL PBS per well) and incubated for 1 min with 1 µg/mL Hoechst 33342 (ThermoFisher, Cat# H3570) to counterstain nuclei. Virus plaques were visualized under LED epifluorescence using a Nikon Eclipse Ts2R microscope (Nikon Instruments, Inc, Melville, NY), and each well was scored positive or negative if at least one virus plaque was visible. Infectivity in TCID_50_ units was calculated according to the Spearman-Kärber method, as described previously (18).

### Plotting of charged clusters within the UL148 ectodomain

The percentage (charged residues/total residues) in each 6-residue window was plotted using a Python script and visualized using Prism 8 software (GraphPad, Inc., San Diego, CA). Eight local maxima were identified at amino acid positions 47-52, 59-62, 79-83, 133-136, 172-177, 222-225, 223-227, 225-230. Because of their close proximity to each other, the final three clusters were regarded as a single charged cluster for the purposes of constructing mutants.

### Expression vectors encoding HA-tagged UL148 harboring charged-cluster-to-alanine mutations

pcDNA3.1+ UL148HA was generated as follows. A PCR product encoding the HA-tagged *UL148* open reading frame was amplified from template plasmid pEF1α UL148HA (18) using primers UL148_reclone_FWD and UL148_reclone_REV. PCR product was digested with BamHI/EcoRI and ligated into BamHI/EcoRI-digested pcDNA3.1+ (Invitrogen) to yield pcDNA3.1+ UL148HA, which was sequence-confirmed using primers CMV_FWD and BGH_REV.

HA-tagged UL148 was subcloned from pcDNA3.1+ into pSP72 as follows. pcDNA3.1+ UL148HA was digested with BamHI/EcoRI. The ∼1kb fragment was agarose-purified and ligated into BamHI/EcoRI-digested pSP72. Overlapping primers (Table 1) were used to amplify the pSP72 UL148HA template and introduce charged-cluster-to-alanine (CCTA) mutations into the *UL148* open reading frame (ORF). PCR products were digested with DpnI for 1 h at 37°C to remove template plasmid before transformation and isolation of carbenicillin-resistant clones, which were confirmed by enzyme digestion and sequencing using primers SP6_FWD and T7_seq_FWD. These mutated *UL148* sequences were then transferred back to pcDNA3.1+ by BamHI/EcoRI digestion and ligation as above.

### Expression vectors encoding HA-tagged chimpanzee CMV UL148 and rhesus CMV Rh159

pcDNA3.1+ Rh159HA was described previously (29). pcDNA3.1+ encoding HA-tagged chimpanzee CMV UL148 was constructed by Gibson assembly of gBlock ChCMV_UL148_FRAG with BamHI/EcoRI-digested pcDNA3.1+ (Invitrogen) and sequence-confirmed using primers CMV_FWD and BGH_REV. The coding sequence for chimpanzee CMV UL148 (Panine herpesvirus 2 strain Heberling, Genbank/ NCBI Reference Sequence: NC_003521.1) was codon-optimized by Integrated DNA Technologies (Coralville, IA), based on *Homo sapiens* codon usage, to facilitate synthesis as a synthetic dsDNA gBlock.

### Inducible expression vectors encoding EGFP fused to UL148 homologs of human, chimpanzee, and rhesus CMV

Inducible lentiviral vectors pOUPc-UL148-eGFP and pOUPc-Rh159-eGFP were described previously (30). pOUPc encoding chimpanzee CMV UL148 fused to eGFP was constructed as follows. Fragment (i) was amplified using primers Fwd_Ch148_OUPgibs and Rev_Ch148_GFPgibs with pcDNA3.1+ ChCMV UL148HA as a template. Fragment (ii) was amplified using primers Fwd_GFP_Ch148gibs and Rev_GFP_allUL148 using pOUPc-UL148-eGFP as a template. Fragments (i) and (ii) were combined with AgeI/MluI-digested pOUPc in Gibson assembly to yield pOUPc-Ch148-eGFP, which was sequence-confirmed using primers CMVcrsnull_Fw, Ubc_Rv, and Ch148_3pr_Fw.

### Construction of recombinant viruses

Synthetic double-stranded DNA (gBlocks) and oligonucleotides were purchased from Integrated DNA Technologies; sequences are provided in **Table 1**. All mutagenesis procedures was performed in the context of the infectious BAC clone of HCMV strain TB40/E, TB40-BAC4 (53), as previously described (17, 18, 29, 52, 54). Briefly, two-step Red “*en passant*” recombination (39, 40) was used in conjunction with GS1783 *E. coli* (a gift of Greg Smith, Northwestern University) harboring derivatives of TB40-BAC4. Recombinant BACs were confirmed by Sanger-sequencing (Genewiz, Inc., South Plainfield, NJ) of modified loci (not shown), and by BamHI and EcoRI restriction enzyme digestion analysis to exclude spurious rearrangements (not shown).

### Recombinant HCMVs encoding HA-tagged UL148 and S-tagged glycoprotein O

Recombinant HCMV, strain TB40/E, encoding S-tagged glycoprotein O and HA-tagged UL148 (TB_148^HA^_gO^S^) was constructed by insertion of a S-tag at the C-terminus of the *UL74* open reading frame, essentially as in (18), but in the context of recombinant TB_148^HA^ (17). Briefly, plasmid pSP72-*UL74*-*S*-*ISceKan* (18) was digested with EcoRV, and the 1.4 kb band was agarose-purified and recombined into TB_148^HA^. Kan-resistant integrate colonies were isolated and resolved of the Kan cassette by L(+)-arabinose induction of the homing endonuclease ISce-I and the bacteriophage lambda Red recombinase system. Kan-sensitive clones were sequence-confirmed using primers TB_UL74_Cterm_seq_FWD and TB_UL74_Cterm_seq_REV.

### Recombinant HCMVs encoding CCTA mutations within *UL148*

CCTA mutations were introduced into TB_148^HA^_gO^S^ as follows. A PCR product containing an I-SceI restriction site upstream of a kanamycin (Kan) resistance cassette and 40-bp flanks homologous to the sequence surrounding each *UL148* charged cluster was amplified using primers indicated in Table 1 and TB_148_ISceKan BAC DNA (17) as the template. The gel-purified PCR product was electroporated into GS1783 *E. coli* carrying an infectious BAC clone of TB_148^HA^_gO^S^. After isolating Kan-resistant colonies, the Kan expression cassette was “scarlessly” resolved by inducing expression of the I-SceI homing endonuclease and Red recombinase. The final Kan-sensitive TB_148^HA^_CCTA_gO^S^ recombinants were sequence-confirmed using primers TB_Rh159HA_seq_FWD and TB_Rh159HA_seq_REV.

### Antibodies

Antibody dilutions for Western blot included 1:1000 rabbit anti-HA (Bethyl), 1:250 mouse anti-CD58 (Invitrogen #MA5800). For flow cytometry, APC-conjugated mouse anti-CD58 antibody (Invitrogen #17-0578) and isotype control (Invitrogen #17-4714) were used at 1:40 dilution. For confocal microscopy, chicken anti-HA (Bethyl) and rabbit anti-Hrd1 (Cell Signaling Technology) were used at a dilution of 1:200. Goat anti-chicken Alexa Fluor 647 (Invitrogen) and goat anti-rabbit Alexa Fluor 488 (Invitrogen) secondary antibodies were used at 1:1000.

### Surface CD58 quantification by flow cytometry

For experiments in transfected cells, 1.2 × 10^6^ A549 cells were electroporated using a Lonza Nucleofector^TM^ 2B device (program X-001) with 1.5 µg pcDNA3.1+ encoding the indicated protein plus 1 µg pmaxGFP^TM^ (Amaxa) in Ingenio nucleofection buffer (Mirus, Inc., cat # MIR 50114). Electroporated cells were then seeded to a 6-well cluster dish and incubated at 37°C until 72 h post-transfection. For experiments in infected fibroblasts, 1.0 × 10^5^ cells were seeded into 6-well cluster dishes and infected at an MOI of 2 TCID_50_/cell with TB_148_HA__gO^S^ or the derived CCTA mutant viruses. The following day, the viral inocula were removed, and media were replenished with fresh 5% newborn calf serum (NCS)/DMEM. The cells were incubated at 37°C until 72 h post-infection. Infected cells were formalin-fixed before the CD58-staining step below.

At time of analysis, cells were detached using a 1:10 dilution of TrypLE^TM^ Express (Gibco #12605) solution. Cells were pelleted (5 min, 300 × *g*) and resuspended in FACS buffer [5% NCS in PBS]. Cells were incubated with APC-conjugated anti-CD58 antibody or isotype control for 1 h at 4°C with rotation, then washed three times with FACS buffer and analyzed by flow cytometry. In transfection experiments, cells were first gated on GFP to remove any non-transfected cells from the analysis. Median APC fluorescence values were determined for each condition, and all values were normalized to the median APC signal of GFP+ cells transfected with empty pcDNA3.1+. Graphs were generated in Prism 8 for Mac Os (GraphPad Software, Inc. San Diego, CA).

### Analysis of protein solubility

2 × 10^5^ fibroblasts (HFFT) were infected at an MOI of 1 TCID_50_ per cell with TB_148^HA^_gO^S^ or the derived CCTA mutants. The following day, cells were washed once with PBS and replenished with fresh 5% NCS/DMEM. At 72 h post-infection, cells were prepared for Western blot analysis of detergent-soluble and - insoluble material as previously described (30). Briefly, cells were washed with PBS and lysed by direct addition of RIPA buffer [25 mM 4-(2-hydroxyethyl)-1-piperazineethanesulfonic acid (HEPES) pH 7.5, 400 mM NaCl, 0.1% sodium dodecyl sulfate (SDS), 0.5% sodium deoxycholate, 1% NP-40], supplemented with 1× protease inhibitor cocktail (ApexBio, Inc., Houston, TX, Cat #K1007) and rotated for 1 h at 4°C. Lysates were centrifuged (21,000 × *g*, 35 min) to separate soluble and insoluble material. The supernatant containing soluble material was transferred to a new tube, and Laemmli buffer was added to 1× final [50 mM Tris (pH 6.8), 2% SDS, 10% glycerol, 0.02% bromophenol blue]. Pellet containing insoluble material was lysed in 1× Laemmli buffer. Samples were reduced by addition of beta-mercaptoethanol (5% final, v/v) and boiled at 90°C for 10 min. Samples were resolved by SDS-PAGE (12% polyacrylamide gel) and transferred to nitrocellulose membrane. Membranes were blotted with indicated antibodies and visualized by Odysey CLx scanner (Li-Cor, Inc., Lincoln, NE).

### Plasmid transfection

For Western blot studies, 1 × 10^6^ HEK-293T (293T) cells were reverse-transfected with 2.5 µg pcDNA3.1+ encoding the indicated proteins, as follows. TransIT-293 transfection reagent (Mirus Inc., Madison, WI, Cat #2700) was mixed with plasmid DNA in Opti-MEM (ThermoFisher) according to the manufacturer’s instructions. After 25 min incubation at room temperature, the transfection reagent-DNA complexes were added to 2 mL 5% FBS/DMEM containing 1 × 10^6^ HEK-293T cells in suspension. Cells and transfection reagent-DNA complexes were gently mixed and seeded to a 6-well plate.

For immunofluorescence imaging, 7 × 10^4^ 293T cells were reverse-transfected with 1 µg doxycycline-inducible pOUPc plasmid (29, 30) encoding the indicated protein as follows. Plasmid DNA was combined with 7.5 µL of 1 mg/mL branched polyethylenimine (PEI; Sigma #408727) in 160 µL of Opti-MEM and gently mixed. DNA-PEI complexes were allowed to form at room temperature for 20 min. 1 mL of 5% FBS/DMEM containing 7 × 10^4^ 293T cells was then added to the DNA-PEI solution and mixed gently before seeding to a 96-well plate. After cells had attached to the well (1 h post-transfection), doxycycline hyclate stock was added directly to each well to a final concentration of 100 ng/mL. Cells were incubated at 37°C for 48 h before imaging for GFP signal on a Nikon Eclipse Ts2R microscope (Nikon Instruments, Inc, Melville, NY).

### Confocal microscopy

7 × 10^5^ fibroblasts (HFFT) were seeded on 12 mm round coverslips (Azer Scientific, #200121) in a 24-well plate. The following day, cells were infected at an MOI of 1 TCID_50_ per cell with TB_148^HA^_gO^S^ or the indicated recombinants. At 72 h post-infection, cells were processed for confocal microscopy essentially as in (30). Briefly, cells were washed once with 1 mL PBS, fixed in 4% (w/v) paraformaldehyde/PBS for 20 min at room temperature, and permeabilized for 3 min by 0.1% TritonX-100. Coverslips were incubated in 5% (v/v) normal goat serum for 30 min at room temperature. After washing three times with 1 mL PBS, coverslips were incubated with 1% Human Fc Block (BD Biosciences, Cat # 564219) for 30 min at room temperature. Coverslips were incubated with primary antibody for 1 h in a humid 37°C incubator, washed three times in 1 mL PBS and then incubated in the presence of secondary antibody (1 h, 37°C humid incubator), followed by three washes in PBS, as above. Finally, coverslips were mounted onto glass slides in mounting medium [0.01 M Tris (pH 8.8), 90% glycerol, 0.5% N-propyl gallate] and sealed with nail polish. Cells were imaged on a Leica TCS SP5 confocal microscope using a Leica 63× oil immersion objective lens.

## ACKNOWLEDGMENTS

This project was supported by NIH Grants R01-AI116851 (to J.P.K) and P30-GM110703. Its contents are solely the responsibility of the authors and do not necessarily represent the official views of the funding agencies. C.C.N. was supported by a Malcolm Feist Predoctoral Fellowship from the Center of Cardiovascular Diseases and Sciences at LSU Health Sciences Center, Shreveport. Author contributions are as follows: C.C.N. conceived of the study, designed research, constructed new recombinant HCMVs and plasmid constructs, carried out most experiments, and contributed to writing the manuscript. A.J.D. performed and contributed to the design of the flow cytometry experiments with infected cells. J.P.K. designed research, interpreted data, obtained funding, and contributed to writing the manuscript. H.Z. assisted with confocal microscopy and constructed the recombinant virus TB_148^HA^_gO^S^.

## Notes

#### Summary of Updates

Typos corrected in main manuscript text and in Figure legends. Minor revisions to Introduction and Discussion sections. References added.

